# Extensive characterization of a Williams Syndrome murine model shows *Gtf2ird1*-mediated rescue of select sensorimotor tasks, but no effect on enhanced social behavior

**DOI:** 10.1101/2023.01.18.523029

**Authors:** Kayla R. Nygaard, Susan E. Maloney, Raylynn G. Swift, Katherine B. McCullough, Rachael E. Wagner, Stuart B. Fass, Krassimira Garbett, Karoly Mirnics, Jeremy Veenstra-VanderWeele, Joseph D. Dougherty

**Affiliations:** Department of Genetics, Washington University School of Medicine, St. Louis, MO 63110, USA; Department of Psychiatry, Washington University School of Medicine, St. Louis, MO 63110, USA; Intellectual & Developmental Disabilities Research Center, Washington University School of Medicine, St. Louis, MO 63110, USA; Department of Pharmacology, Vanderbilt University; Psychiatry, Biochemistry & Molecular Biology, Pharmacology & Experimental Neuroscience, Munroe-Meyer Institute for Genetics and Rehabilitation, University of Nebraska Medical Center Omaha, NE 68198-5450; Departments of Psychiatry and Pediatrics, Columbia University; New York State Psychiatric Institute; and Center for Autism and the Developing Brain, New York-Presbyterian Hospital

**Author notes:** Corresponding author Dr. Joseph D. Dougherty, Professor, Department of Psychiatry, Department of Genetics, Washington University School of Medicine, 660 South Euclid Avenue, St. Louis, MO 63110-1093.

**Keywords:** Williams Syndrome, Gtf2ird1, anxiety, sociability, motor function, mouse model

## Abstract

Williams Syndrome is a rare neurodevelopmental disorder exhibiting cognitive and behavioral abnormalities, including increased social motivation, risk of anxiety and specific phobias along with perturbed motor function. Williams Syndrome is caused by a microdeletion of 26-28 genes on chromosome 7, including *GTF2IRD1*, which encodes a transcription factor suggested to play a role in the behavioral profile of Williams Syndrome. Duplications of the full region also lead to frequent autism diagnosis, social phobias, and language delay. Thus, genes in the region appear to regulate social motivation in a dose-sensitive manner. A ‘Complete Deletion’ mouse, heterozygously eliminating the syntenic Williams Syndrome region, has been deeply characterized for cardiac phenotypes, but direct measures of social motivation have not been assessed. Furthermore, the role of *Gtf2ird1* in these behaviors has not been addressed in a relevant genetic context. Here, we have generated a mouse overexpressing *Gtf2ird1*, which can be used both to model duplication of this gene alone and to rescue *Gtf2ird1* expression in the Complete Deletion mice. Using a comprehensive behavioral pipeline and direct measures of social motivation, we provide evidence that the Williams Syndrome Critical Region regulates social motivation along with motor and anxiety phenotypes, but that *Gtf2ird1* complementation is not sufficient to rescue most of these traits, and duplication does not decrease social motivation. However, *Gtf2ird1* complementation does rescue light-aversive behavior and performance on select sensorimotor tasks, perhaps indicating a role for this gene in sensory processing or integration.

## Introduction

Microdeletion of the 7q11.23 region results in the neurodevelopmental disorder known as Williams Syndrome (WS). Hemizygosity of the 26-28 genes in this region, also known as the Williams Syndrome Critical Region (WSCR), causes multisystemic symptoms which match some features and mirror others from the reciprocal 7q11.23 Duplication Syndrome (Dup 7), revealing the importance of gene dosage in the pathophysiology of these disorders (Adamo *et al*. 2014; Strong *et al*. 2015; Zanella *et al*. 2019). Both syndromes result in altered craniofacial features, cardiac issues, motor coordination deficits, and behavioral challenges (Morris *et al*. 2015; Kozel *et al*. 2021). Many people with WS exhibit hypersociability and tend to approach strangers with little apprehension, though the lack of social anxiety does not preclude a more generalized anxiety and occasional extreme phobias, both of which are more prevalent in individuals with WS than the general population (Doyle *et al*. 2004; Levitin *et al*. 2005; Berg *et al*. 2007). Unfortunately, the underlying mechanisms of these behavioral differences are not well understood and thus no targeted treatments exist to help individuals with WS navigate the expectations of society, similar to the struggle autistic individuals face. However, unlike the complex etiology of idiopathic autism, the discrete genetic foundation of WS provides a unique opportunity to uncover these mechanisms, as a relatively small deletion leads to such a recognizable behavioral profile.

As WS and Dup7 are rare, understanding the complex etiology and circuit pathology underlying behavioral phenotypes in humans, or with human brain samples, is challenging. While cellular phenotypes can be investigated in iPSC models (Adamo *et al*. 2014), animal models are still required to uncover the link between gene dosage and behavioral phenotypes. Fortunately, the WSCR is syntenic in mice, and a complete deletion (CD) mouse model eliminating most of the region has been developed that recapitulates many features of WS (Segura-Puimedon *et al*. 2014).

An initial survey of various features in the CD mouse line discovered mild cardiac deficits, craniofacial anomalies, and some alterations in behavior. Specifically, the authors reported deficits in motor performance in a Rotarod task and a decreased habituation to social stimuli in an open field social interaction test (Segura-Puimedon *et al*. 2014). However, the previous measures on pure C57BL/6J showing increased social interest were only conducted in males and they did not utilize the classic 3-Chamber Social Approach task. In our hands, the three-Chamber Social Approach task showed no difference in CD mice (albeit on a FVB/AntJ x C57BL/6J hybrid background) (Kopp *et al*. 2019); the FVB strain generally shows less social approach than C57BL/6J (Nygaard *et al*. 2019), suggesting this background may be less sensitive for CD social phenotyping. Furthermore, social motivation, the amount of work an animal is willing to do to engage with a conspecific, has not been directly measured. Likewise, while motor learning has been assessed on the Rotarod apparatus (Segura-Puimedon *et al*. 2014), less work has characterized motor strength or coordination generally and the results were not consistent on the hybrid background with slightly different test parameters (Kopp *et al*. 2019). Finally, anxiety has also been a difficult domain to assess consistently in mice, as transient emotionality can affect the results (Ramos 2008). For example, similar mouse models deleting a single gene in the WSCR, *Gtf2ird1*, show opposite results in anxiety-like behavior (Young *et al*. 2008; Schneider *et al*. 2012). Overall, a deeper phenotyping of these domains would be of use, especially to provide a foundation for studies that address the effects of WSCR copy number variation at the level of mechanisms or circuits.

Prior to the development of the CD line modeling the full deletion, single gene deletions were the most common approach in trying to elucidate function. *Gtf2ird1* is one such gene that has been implicated in a variety of hallmark WS phenotypes, from craniofacial to cognitive and behavioral differences. *Gtf2ird1* is often implicated alongside its neighbor and family member, *Gtf2i*. As these genes occur in tandem in the WSCR and are rarely found separately affected by atypical deletions, it is difficult to isolate their effects using human studies alone. While both genes are conserved in the mouse genome, there has been more difficulty with reliably producing a *Gtf2ird1* knockout animal. Alternative and frame-shifted start codons allow truncated versions of the protein to be expressed, even preserving much of its function outside of its negative autoregulation (Kopp *et al*. 2020). Verifying *Gtf2ird1* expression, or lack thereof, was also unreliable prior to the relatively recent development of effective antibodies.

To avoid the trouble of deleting this elusive gene, we adopted a different strategy to gauge the influence of *Gtf2ird1* on relevant phenotypes; we designed a study to assess the impact of *Gtf2ird1* while also providing an extensive characterization of the CD model, as both of these contributions would benefit our understanding of the WSCR. Thus, we present a novel *Gtf2ird1* transgenic expression line, which we use to thoroughly assess the role of *Gtf2ird1*. We test the hypothesis that *Gtf2ird1* plays a dose-dependent role in the cognitive and behavioral symptoms of WS and concurrently examined the effects of *Gtf2ird1* overexpression on the C57BL/6J pure background and in the presence of WSCR deletion. We used a comprehensive battery of tasks designed to elucidate the contributions of *Gtf2ird1* to WS-relevant phenotypes. Simultaneously, using the same extensive suite of behavioral measures, we provide a detailed assessment of the CD mouse, providing key information on additional phenotypes related to motor, anxiety, fear, and social behaviors, broadening the initial characterization.

In these studies, we replicate and extend the previously reported social differences in the CD mice (Segura-Puimedon *et al*. 2014), showing enhanced social approach and motivation, in addition to sensorimotor differences and greater avoidance behavior in some anxiety-related tasks. Finally, we rule out *Gtf2ird1* as being the sole mediator of the social changes, as duplication of this gene did not decrease these behaviors, nor did its complementation of the complete deletion rescue any notable social phenotypes. However, it does appear to mediate aspects of light-induced anxiety-related behaviors and sensorimotor coordination, as complementation can ameliorate the deficits observed in the CD mice, suggesting a role for *Gtf2ird1* in sensorimotor processing.

## Materials and Methods

All experimental protocols were approved by and performed in accordance with the relevant guidelines and regulations of the Institutional Animal Care and Use Committee of Washington University in St. Louis and were in compliance with US National Research Council’s Guide for the Care and Use of Laboratory Animals, the US Public Health Service’s Policy on Humane Care and Use of Laboratory Animals, and Guide for the Care and Use of Laboratory Animals.

### *Gtf2ird1* Transgenic Mouse Creation

We selected a bacterial artificial chromosome (BAC) clone (RP24-508D22) which contained the entirety of the 100 kb *Gtf2ird1* gene, and 89 kb (60 kb at the 5’ end and 28 kb at the 3’) of flanking regulatory sequence (e.g., the *Gtf2ird1* promoter, etc.), but no additional intact genes or their promoters. This was then recombineered using standard methods to insert an HA tag in-frame directly before the stop codon of the beta isoforms (Tay *et al*. 2003; Ebert *et al*. 2006). Specifically, we used homologous recombination via transient expression of RecA, followed by Neomycin selection of an inserted FRT flanked cassette. The selection cassette was then removed after the expression of the FLPe recombinase, leaving behind a single FRT site downstream of the stop codon. Transgenic mice overexpressing *Gtf2ird1* (TG-Gtf2ird1-HA or simply TG) were created by injecting this modified BAC into C57BL/6NTac mouse oocytes and transplanting these eggs into pseudopregnant surrogates to carry them to term. Transgene-specific primers (BgenoF3 – CAACATTCCCAAGCGCAAGAG and BgenoR3 – GATAACTGATCGCGGCCAGC, which produce a 440 bp product in TG animals and no product in WT animals) were used for identification of TG founder animals. BAC copy number was determined to be 2-4. Multiple founders were evaluated to confirm transgenic RNA production by RT-PCR, and a single line was taken forward for evaluation. Lines were backcrossed to C57BL6/J for over four generations prior to commencing experiments.

### Husbandry

All mice used in this study were maintained and bred in the vivarium at Washington University in St. Louis on a 12/12 hour light/dark cycle with food and water provided freely. Three distinct mouse lines were used: C57BL/6J (WT; RRID:IMSR_JAX:000664), the Complete Deletion (CD) mouse modeling deletion of the Williams Syndrome critical region (Segura-Puimedon *et al*. 2014), and a novel transgenic line (TG) overexpressing *Gtf2ird1* with an HA tag. CD and TG lines were maintained as heterozygotes by crossing to WT animals. Heterozygous CD and TG mice were crossed to produce behavioral cohorts containing WT, TG, CD, and TG/CD littermates to best compare across genotypes. Animals were housed by genotype and sex at weaning. Tissue was collected from pups for initial genotyping and again after death to verify genotype via PCR amplification.

### Molecular Validation

Molecular analysis to assess RNA and protein levels via RT-qPCR and Western blotting was performed as previously described (Kopp *et al*. 2020). Brains were collected from pups ∼E13.5 for initial characterization of the novel line and just prior to weaning at postnatal days 19 or 20 (P19-P20) for validation of the crosses. RNA expression and protein levels were assessed relative to *Gapdh* using primers and antibodies described previously (Kopp *et al*. 2020). Expression of the transgenic allele was verified by a Western blot using an antibody to the HA tag, which is included only on some transcripts of *Gtf2ird1* due to alternative splicing of the last exon.

### Behavioral Testing

For behavioral analysis, three separate cohorts of mice were used to assess a variety of characteristics (**Table 1**). All tasks were run by female experimenters. Tasks within each behavioral battery were ordered from least to most stressful to minimize the effect one task had on subsequent tasks. Adolescent and adult mice were handled for 5 days prior to starting the first behavioral task and the tails of mice in Cohorts 1 and 2 were marked with a non-toxic, permanent marker during weight collection and regularly thereafter to easily distinguish mice during testing. Males were run before female animals to avoid olfactory cue influence on behavior. Testing orders were randomly counterbalanced for group across apparatuses and trials. Unless otherwise indicated, all equipment was cleaned between animals with a 0.02% chlorhexidine diacetate solution (Nolvasan, Zoetis).

**Table 1.** Behavioral cohort sample size and task order.

| COHORT 1 |  | COHORT 2 |  | COHORT 3 |  |
| --- | --- | --- | --- | --- | --- |
| Total N=94 (48F, 46M) |  | Total N=77 (40F, 37M) |  | Total N=91 (49F, 42M) |  |
| WT=29 TG=21,<br>CD=20 TG/CD=24 |  | WT=20 TG=21<br>CD=17 TG/CD=19 |  | WT=30 TG=15<br>CD=24 TG/CD=22 |  |
| TASK | Age | TASK | Age | TASK | Age |
| Open Field | P31<br>(27-36) | Sensorimotor Battery | P75<br>(56-105) | Temp/Weight | P5,7,9 |
| Social Approach | P32<br>(28-37) | Rotarod | P78<br>(59-108) | USVs | P5,7,9 |
| Marble Burying | P36<br>(30-40) | Light/Dark Box | P79<br>(60-109) | Righting Reflex | P14 |
| Elevated Plus Maze | P39<br>(33-43) | 3-Chamber Social Approach | P83<br>(64-113) |  |  |
| Novel Object | P44<br>(38-47) | Tube Test | P89<br>(70-119) |  |  |
| Social Operant | P56<br>(40-97) | PrePulse Inhibition | P96<br>(77-126) |  |  |
| Conditioned Fear | P78<br>(59-126) | Resident Intruder | P108<br>(89-138) |  |  |

*\*Age is the average postnatal day for all mice with the full age range provided below.*

### COHORT 1

#### Open Field

To assess activity levels and passive avoidance behavior, we used the open field task adapted from our previously published methods (Chen *et al*. 2021). Briefly, mice were placed in a 50 × 50 × 45 cm clear acrylic enclosure under red light at 9 lux, within a sound- and scent-attenuated white opaque box (70.5 × 50.5 × 60 cm) to minimize external stimuli, and allowed to freely explore for 60 minutes. Any-Maze software (Stoelting, Co) tracked the movement via the body center, beginning when the doors to the chamber were closed via video captured with an overhead CCTV camera. A center zone was designated as the middle 50% of the total chamber area. Movement of the animal throughout the arena was quantified as distance traveled, and as time spent in and entries into the center and perimeter zones. The center zone was designated as the inner 50% of the open field area, and the perimeter was the outer 50%.

#### Open Field Social Approach

Replicating previously published methods (Sakurai *et al*. 2011; Segura-Puimedon *et al*. 2014), we examined social approach behavior in an open field setting, under white light at 50 lux. In the center of the Open Field enclosure described above, a novel, sex- and age-matched stimulus animal (C57BL/6J) was placed under a wire pencil cup (Galaxy Pencil/Utility Cup, Spectrum Diversified), with a clear plastic cup on top to prevent climbing. The experimental animal was then added to the chamber and allowed to explore and interact with the social stimulus. After 15 minutes, recording was stopped, the experimental animal was removed to a clean holding chamber, and the stimulus animal was switched out with a novel mouse. The experimental animal was returned to the chamber for 5 additional minutes of recorded exploration. Any-Maze video tracking was used for both trials, and measured distance traveled and time spent in an investigation zone defined as 2 cm around the circumference of the cup. Wire cups were cleaned with 70% ethanol between mice.

### Elevated Plus Maze

Anxiety-like behaviors were tested using the Elevated Plus Maze (EPM) as previously described (Kopp *et al*. 2019). Briefly, mice were placed in the center of the apparatus, which contained two open and two closed arms, and allowed to explore for 5 minutes in the dark. This was repeated for two more days. Trials were recorded under infrared illumination with an overhead camera using Ethovision software (Noldus Information Technology) to track movement of the animal in the apparatus.

### Marble Burying

The Marble Burying task was used to assess compulsive digging behavior, adapted from previous methods (Maloney *et al*. 2018a). As in prior work, the mice were introduced to a novel, transparent enclosure (47.6 × 25.4 × 20.6 cm, contained within a sound- and scent-attenuated white opaque box (70.5 × 50.5 × 60 cm) to minimize external stimuli) with 20 evenly spaced, clear marbles on clean, novel, autoclaved aspen bedding. Animals were allowed to explore freely for 30 minutes. After the animals were removed, two independent scorers recorded the number of marbles buried (defined as at least two-thirds covered with bedding). These scores were averaged for analysis. Between mice, the marbles were cleaned with 70% ethanol. For this study, we also tracked the animals’ movement in the apparatus via Any-maze tracking software and quantified distance traveled for overall activity levels and time spent in the center 50% of the arena.

### Open Field Novel Object Exploration

Adapting previously published methods (Segura-Puimedon *et al*. 2014), we examined novel object exploration in an open field setting to control for potential novel effects in the Open Field Social Approach task. A translucent cube was placed in the center of the same Open Field chamber used for Open Field and Open Field Social Approach described above under white light at 50 lux. Mice were placed in the apparatus to explore freely for 20 minutes while movement was recorded and tracked using Any-Maze software. A 2 cm investigation zone was defined around the object, in addition to center and perimeter areas of the arena.

### Social Motivation Operant Conditioning

Social motivation, or how hard an animal will work for access to a social partner, was assessed using our social motivation operant assay. Our 16-day paradigm allows for assessment of both social reward seeking and social orienting, two components of social motivation (Chevallier *et al*. 2012). We followed the procedure outlined in our previous work (Chen *et al*. 2021; Maloney *et al*. 2022). Briefly, an operant conditioning chamber was modified to include a door that raised in response to a nosepoke in the active hole to provide 12 seconds of access to a novel sex- and age-matched partner stimulus mouse. To assess social reward seeking, the number of active (i.e., elicits a reward) versus inactive nosepokes were quantified. To assess social orienting, the behavior of the animal was tracked using Ethovision (Noldus) and number of interactions with the stimulus mouse and time spent near the stimulus mouse were quantified. Following two days of habituation (door remained open and the nosepoke holes were not accessible), mice received at least 3 days of fixed ratio 1 (FR1) conditioning, where 1 nosepoke in the active hole resulted in a reward. Mice that had at least 40 active nosepokes, 75% accuracy (active:inactive), and 65% successful rewards (interactions during rewards) were considered to have met conditioning criteria and progressed to a fixed ratio of 3 (FR3), where 3 nosepokes were required to receive the reward. After 3 days of FR3 (or 10 days of FR1 for mice who failed to reach criteria), mice were tested in a progressive ratio of 3 (PR3), where the first reward was provided after 3 active nosepokes and each subsequent reward required 3 additional nosepokes to obtain. The breakpoint was measured as the number of rewards a mouse was able to acquire before 30 minutes of nosepoke inactivity.

### Conditioned Fear

To assess associative and anxiety-related memory, mice were tested in a Conditioned Fear task over three days following previously published methods (Maloney *et al*. 2019; Kopp *et al*. 2020; Chen *et al*. 2021). Briefly, following pairing of a tone and context with a 1.0 mA footshock on day 1, all mice were tested for contextual fear memory on day 2 and cued fear memory in response to the tone only on day 3. Shock sensitivity was evaluated as we previously described (Kopp *et al*. 2020).

### COHORT 2

#### Sensorimotor Battery

Adult mice were evaluated with a battery of sensorimotor measures to assess motor initiation, balance, strength, and coordination using previously published methods (Kopp *et al*. 2020; Chen *et al*. 2021). The battery included evaluation of walk initiation, balance (Ledge and Platform tests), fine motor coordination (Pole test), and strength with coordination (Inclined and Inverted Screen tests).

#### Rotarod

Motor coordination was assessed using the Rotarod following our previously published methods (Maloney *et al*. 2019). Briefly, latency to fall was measured for each mouse in three different situations: a stationary rod (for up to 60 sec), a continuously rotating rod (3.0 rpm; for up to 60 sec), and an accelerating rod (3.0-17 rpm; for up to 180 seconds).

### Light/Dark Box

The Light/Dark Box was used to assess anxiety-related passive avoidance behavior leveraging the mouse’s innate preference for dark spaces. Mice were placed in the dark side of a chamber (47.6 × 25.4 × 20.6 cm) and were allowed to explore freely. The light side, which was twice as large as the dark side, was illuminated at 65 lux with incandescent desk lamps. Beam brakes were used to measure time spent in each chamber during the 6-minute task. For the first two minutes, mice were confined to the dark side of the apparatus, then mice had four minutes to explore the entire apparatus. Time spent in and latency to move to the light side during the latter four minutes was used as a proxy for anxiety-like behavior, with more anxious-like mice avoiding the brightly lit open space.

### 3-Chamber Social Approach

Sociability and preference for social novelty were examined in the Social Approach task, following our previously published methods (Chen *et al*. 2021). The mice received two, 10-min habituation trials: first to the center chamber of the apparatus and then to the entire chamber including the empty social investigation cups. Next, sociability was assessed for 10 min during which a novel age- and sex-matched conspecific was placed under one cup (the side used was counterbalanced across groups). During the fourth 10-min trial, a second, age- and sex-matched novel conspecific was placed under the other cup to assess preference for social novelty. The time spent investigating and number of investigations for each investigation cup, as well as time in and entries into each chamber and total distance traveled, was quantified using Any-Maze video tracking software.

### Tube Test of Social Dominance

As social creatures, mice create social hierarchies within their social groups. Thus, laboratory mice acquire social hierarchical rank behaviors within their cage environments between six-eight weeks of age, which can be leveraged to examine normal social dominance behavior. We tested for this normal hierarchical behavior in our mice using the tube test for social dominance following our previously described methods (Maloney *et al*. 2018a; Chen *et al*. 2021).

### Acoustic Startle/Pre-Pulse Inhibition Task

Sensorimotor gating and startle reactivity were assessed using the Acoustic Startle/Pre-Pulse Inhibition (PPI) task following our previously published methods (Kopp *et al*. 2020; Chen *et al*. 2021). Briefly, acoustic startle to a 120 dB auditory stimulus pulse (40 ms broadband burst) and PPI (response to a pre-pulse plus the startle pulse) were measured concurrently using computerized instrumentation (StartleMonitor, Kinder Scientific) over 65 randomized trials. A percent PPI score for each trial was calculated using the following equation: % PPI = (startle pulse alone − (pre-pulse + startle pulse))/startle pulse alone × 100.

### Resident Intruder

To assess agnostic, and thus aggression-related, behaviors, we used the Resident Intruder paradigm that leverages the propensity of male mice to defend their territory from an unfamiliar male as previously described (Yuede *et al*. 2013; Kopp *et al*. 2019). Briefly, male mice were single housed for 6 days followed by 4 more days co-housed with a female to establish a territory. Twenty-four hours prior to testing, the female was removed from the cages. Across three test days, the male cages were placed in a sound- and scent-attenuated white opaque box (70.5 × 50.5 × 60 cm) to minimize external stimuli). On each test day, a novel juvenile CD1 male mouse served as the intruder and was placed into the test animal’s home cage for 10 min, which was video recorded.

For analysis, we used the DeepLabCut (DLC) neural network for pose estimation, version 2.2rc3 using the resnetv50 (Nath *et al*. 2019), followed by the Simple Behavior Annotator (simBA) version simba-uw-tf 0.85.3 (Nilsson *et al*. 2020) random forest classifier generated from the pose estimates for classification of attack behaviors. Specifically, we labeled 240 frames taken from 120 approximately ten-minute videos that were converted from MTS to mp4 using ffmpeg. Each frame was labeled with sixteen unique body parts, eight per animal as according to the simBA 16bp user manual. The DLC neural net trained using 80% of the labeled frames for approximately 370,000 iterations with default parameters. With a p-cutoff of 0.9, the trained network was able to predict high confidence mouse body part positions within 2.26 millimeters of human-labeled positions in the testing set, and the general quality of labels were confirmed by visual inspection of several videos. Estimates of pose were then exported to .csv for analysis by simBA. SimBA was trained to identify attack behavior (resident attacking intruder, RI) using 180 annotated behavior files, downloaded from https://osf.io/tmu6y/ in addition to four in-house annotated videos. All training files were annotated according to definitions found in the simBA preprint (Nilsson *et al*. 2020).

In addition to these RI annotated files, a custom script was created to reverse the direction of attack in order to estimate instances of the intruder attacking the resident (IR). Both training sets were trained using 6000 trees, 20% training set, Gini impurity function, number of estimators equal to the squared number of features, and 1 min leaf. Probability thresholds for each model were chosen based on a maximum F1 score curve from the testing set. To ensure IR and RI datasets were mutually exclusive, a custom script was written to calculate the mean of random forest probability of overlapping frames of RI and IR behaviors and keep only the behavior with the larger mean probability across the overlap. Scoring by algorithms was visually inspected by trained behaviorists for a subset of videos to confirm accuracy. All custom code is available upon reasonable request.

### COHORT 3

#### Maternal separation induced ultrasonic vocalizations and righting reflex

We assessed the developmental trajectory of early postnatal ultrasonic vocalizations (USVs) and acquisition of physical and reflex milestones as previously described (Maloney *et al*. 2018b; Chen *et al*. 2021). Briefly, pups were tested in their colony room by the same female experimenter on P5, 7, and 9. Mice were identified and genotyped by toe clipping, which was performed after the P5 recording session. All recordings occurred after 12 PM CST between March and September of the same year. Prior to recording, the parents were removed, and pups in the nest were placed in a warming chamber at 33°C without removing them from their nest to maintain a surface body temperature of 31.1–37.5°C. After 10 minutes to acclimate body temperature, each pup was placed in an empty cage in a sound attenuating box (36×64×60 cm) and recorded for 3 minutes. The Avisoft UltraSoundGate CM16 microphone was positioned 5 cm from the top of the cage and an Avisoft UltraSoundGate 116H amplifier (gain = 8, 16 bits, sample rate = 250 kHz) was used for all measurements. USVs were recorded using the Avisoft-RECORDER software. Raw WAV files were processed using a custom MATLAB pipeline to extract call numbers and spectral and temporal call features (Maloney *et al*. 2018b; Chen *et al*. 2021).

In addition to USVs, weight and temperature were recorded for each mouse at each time point. A non-contact HDE Infrared Thermometer was used to take the temperature of each mouse before they were removed from the nest for USV recording. Mice were weighed after recording. Pinnae detachment was also assessed at P5, and eye opening was documented at P14. At P14, the righting reflex was evaluated for each mouse by measuring the time for pup to right itself after being held on its back for 5 seconds as described previously (Chen *et al*. 2021). Three trials, limited to 1 minute, were performed for each mouse, averaged for analysis, and direction of righting was noted.

#### Data Analysis

Analyses were conducted in SPSS v27. Prior to analyses, data was screened for missing values and fit of distributions with assumptions underlying univariate analysis. This included the Shapiro-Wilk test on z-score-transformed data and qq plot investigations for normality, Levene’s test for homogeneity of variance, and boxplot and z-score (±3.29) investigation for identification of influential outliers. Means and standard errors were computed for each measure. All variables were examined via 3-way ANOVA with Bonferroni correction to assess the effects of the CD and TG alleles with sex included as a predictor. If appropriate, sex was dropped from the model to achieve best fit. Weight was used as a covariate for data analysis in the Acoustic Startle/Pre-Pulse Inhibition Task. For tasks with multiple timepoints measured per animal, a repeated measures ANOVA was applied if no data points were missing, otherwise a linear mixed model was used with the repeated predictor included as a random factor nested with subject to create hierarchy. To achieve normality for a given variable, outliers with a z-score ± 3.29 were removed or a square root transformation was applied (conditioned fear data, resident intruder data, social operant data). If normality could not be achieved and/or variance was not homogenous, nonparametric analysis was performed. All graphed data represents the raw values and the standard error of the mean. Details for all statistical tests and results can be found in Supplemental Tables S1 - S4.

## Results

### A novel overexpression mouse rescues *Gtf2ird1* expression in the context of a complete deletion of the syntenic Williams Syndrome Critical Region

Evidence from atypical deletions show the telomeric end of the Williams Syndrome Critical Region (WSCR) is important for most of the key WS features. Two specific genes, *Gtf2i* and *Gtf2ird1*, within the telomeric end are suspected to play important roles in the cognitive and behavioral profiles of WS. While a *Gtf2i* mouse line has been developed, no such line exists for *Gtf2ird1*. Here we fill that gap with a novel transgenic line that expresses *Gtf2ird1* and test the hypothesis that *Gtf2ird1* is critical for features of WS by rescuing its expression on the most relevant background, the complete deletion of the WSCR.

To determine the role *Gtf2ird1* plays in the WS behavioral repertoire, we first generated and validated a novel mouse overexpressing the general transcription factor GTF2IRD1 (TG-Gtf2ird1-HA) via its endogenous regulatory elements, engineered using a bacterial artificial chromosome with an HA tag that was inserted just prior to a stop codon of *Gtf2ird1* (**Fig 1A**). The line was validated through qPCR and Western blot analysis of heterozygous animals which reveal increased production of *Gtf2ird1* RNA (**Fig 1B**; t=-5.247, p=7.76×10^−4^) and protein (**Fig 1C**; t=-1.991, p=0.048, one-tailed).

**Figure 1.**
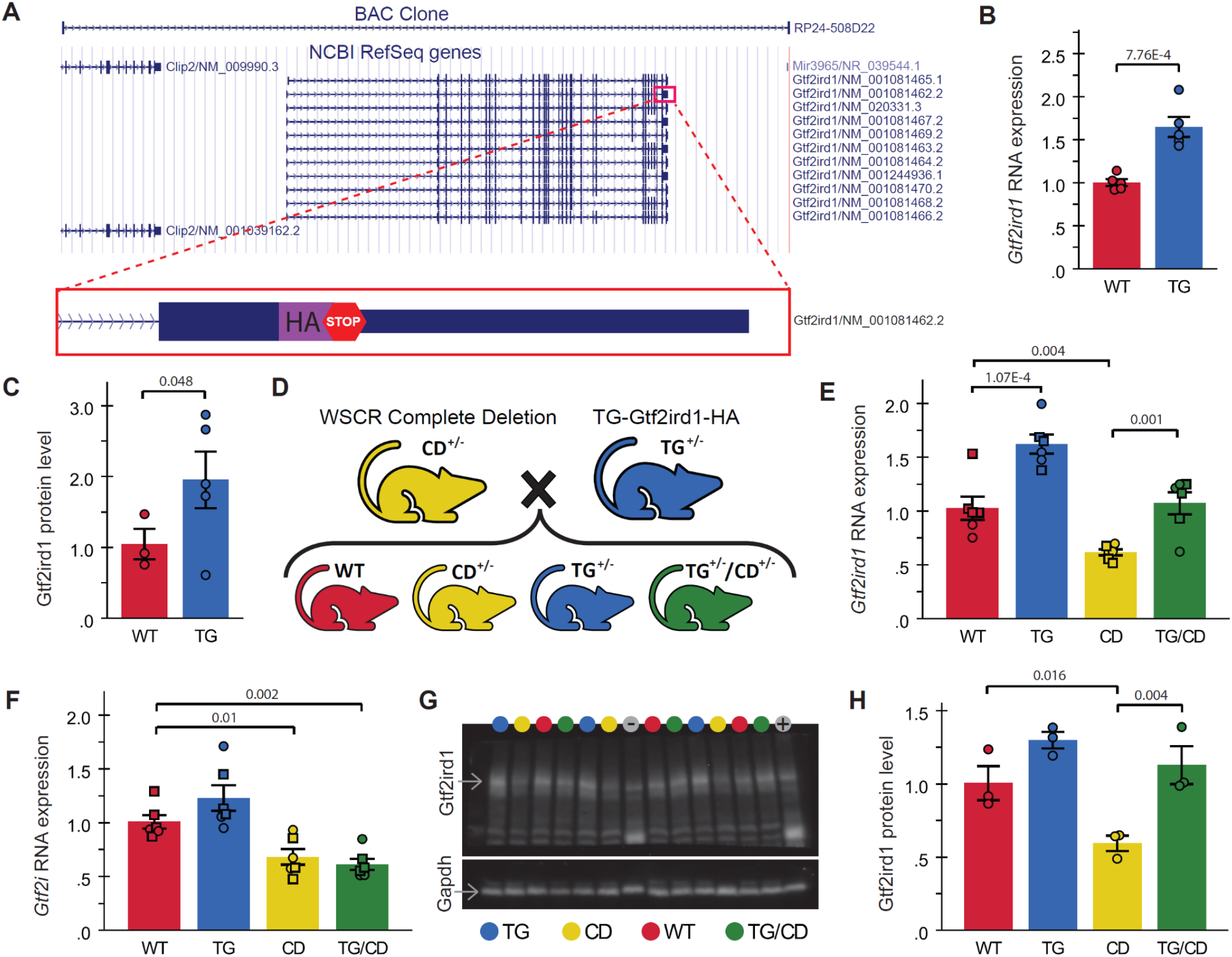
Novel *Gtf2ird1* overexpression mouse rescues *Gtf2ird1* RNA and protein levels in a Complete Deletion mouse modeling deletion of the syntenic Williams Syndrome Critical Region. **A)** Location of the BAC clone used to create the TG-Gtf2ird1-HA mouse line along with a cartoon of the HA-tag added just prior to the stop codon of one of the *Gtf2ird1* isoforms. **B)** RNA expression of *Gtf2ird1* relative to *Gapdh* in WT and TG littermates. **C)** Relative GTF2IRD1 protein levels in WT and TG littermates, n=3 WT, 5 TG. **D)** Heterozygous (+/-) CD and TG animals were crossed to directly compare WT, TG, CD, and TG/CD progeny. **E)** *Gtf2ird1* RNA expression in progeny from cross outlined in panel D, n=6 per genotype. **F)** *Gtf2i* RNA expression from the same animals as in E. **G)** Western blot of TG x CD progeny probed with antibodies for GTF2IRD1 and GAPDH, colored circles above the lanes indicate genotype, - and + represent negative and positive controls for the transgene. **H)** GTF2IRD1 protein levels quantified from the blot in panel G, n=3 per genotype. All RNA and protein levels were normalized to *Gapdh* expression. For E and F only, square = male, circle = female. All statistical details including sample sizes are reported in Table S1.

Next, we demonstrated the ability of the TG-Gtf2ird1-HA mouse to rescue *Gtf2ird1* expression in the Complete Deletion (CD) mouse, a line that effectively deletes the syntenic Williams Syndrome Critical Region (Segura-Puimedon *et al*. 2014), by crossing heterozygous TG-Gtf2ird1-HA and CD animals to produce four distinct progeny: wildtype (WT), TG-Gtf2ird1-HA (TG), Complete Deletion (CD), and the putative rescue (TG/CD), which combines the transgene and the complete deletion (**Fig 1D**). Molecular validation via qPCR confirmed *Gtf2ird1* overexpression in TG heterozygotes and decreased expression in CD animals relative to WTs, while RNA expression in the TG/CD group was not significantly different from WT, indicative of a molecular rescue (**Fig 1E**; F(3,20)=22.190, p=1×10^−6^). To ensure altered expression was specific to *Gtf2ird1*, we also measured relative expression of the nearby related gene, *Gtf2i*. The overexpression of *Gtf2ird1* did not significantly alter RNA expression of *Gtf2i*, which was significantly lower on the CD background regardless of transgene presence (**Fig 1F**; F(3,20)=12.818, p=6.8×10^−5^). To confirm protein expression was also affected, we ran a Western blot, probing for GTF2IRD1 using GAPDH as a control (**Fig 1G**). CD GTF2IRD1 protein levels were significantly lower than WT, TG, and TG/CD GTF2IRD1 levels, which did not differ from each other significantly (**Fig 1H**; F(3,8)=9.918, p=0.005).

Having thus validated the expression of the *Gt2ird1* allele and complementation of *Gtf2ird1* levels in the CD background, we then utilized this same breeding scheme to generate a set of litter-matched behavioral cohorts for comprehensive behavioral testing (**Table 1**), enabling a study of main effects of each allele, as well as detection of interactions. We likewise included sex in all subsequent analyses, and report sex effects when significant.

### *Gtf2ird1* restoration ameliorates select sensorimotor coordination deficits in Complete Deletion mice

Both WS and Dup7 are associated with strength deficits and motor delays. While G*tf2ird1* has been connected to the WS craniofacial phenotype and is suspected to play a role in the unique cognitive profile (which includes visuospatial processing deficits) and behavioral features of WS, its role in sensorimotor features of WS has not been thoroughly defined. To address both the impact of *Gtf2ird1* on these features and the complete WS deletion in CD mice, we devised a comprehensive assessment of sensorimotor abilities, which also provided information necessary to properly interpret tasks relying on adequate motor performance. The wide-ranging compilation of tasks addressed a variety of basic motor abilities and more complex tasks requiring integration of sensory information (in mice, coordinated movement often is informed by their whiskers, rather than their eyes) (Haridas *et al*. 2018). We split relevant tasks across two cohorts; in the first cohort (**Fig 2A**, *above midline*), we tested activity over one hour in an open field apparatus and natural digging behaviors as observed in the Marble Burying task. Animals in the second cohort (**Fig 2A**, *below midline*) were tested using the Rotarod task, Acoustic Startle Response/Pre-Pulse inhibition assay, and a sensorimotor battery, which included Walk, Inverted Screen, Pole, Platform, and Ledge tasks to assess a variety of movement related abilities, such as motor initiation, strength, coordination, and balance, as well as sensory processing.

**Figure 2.**
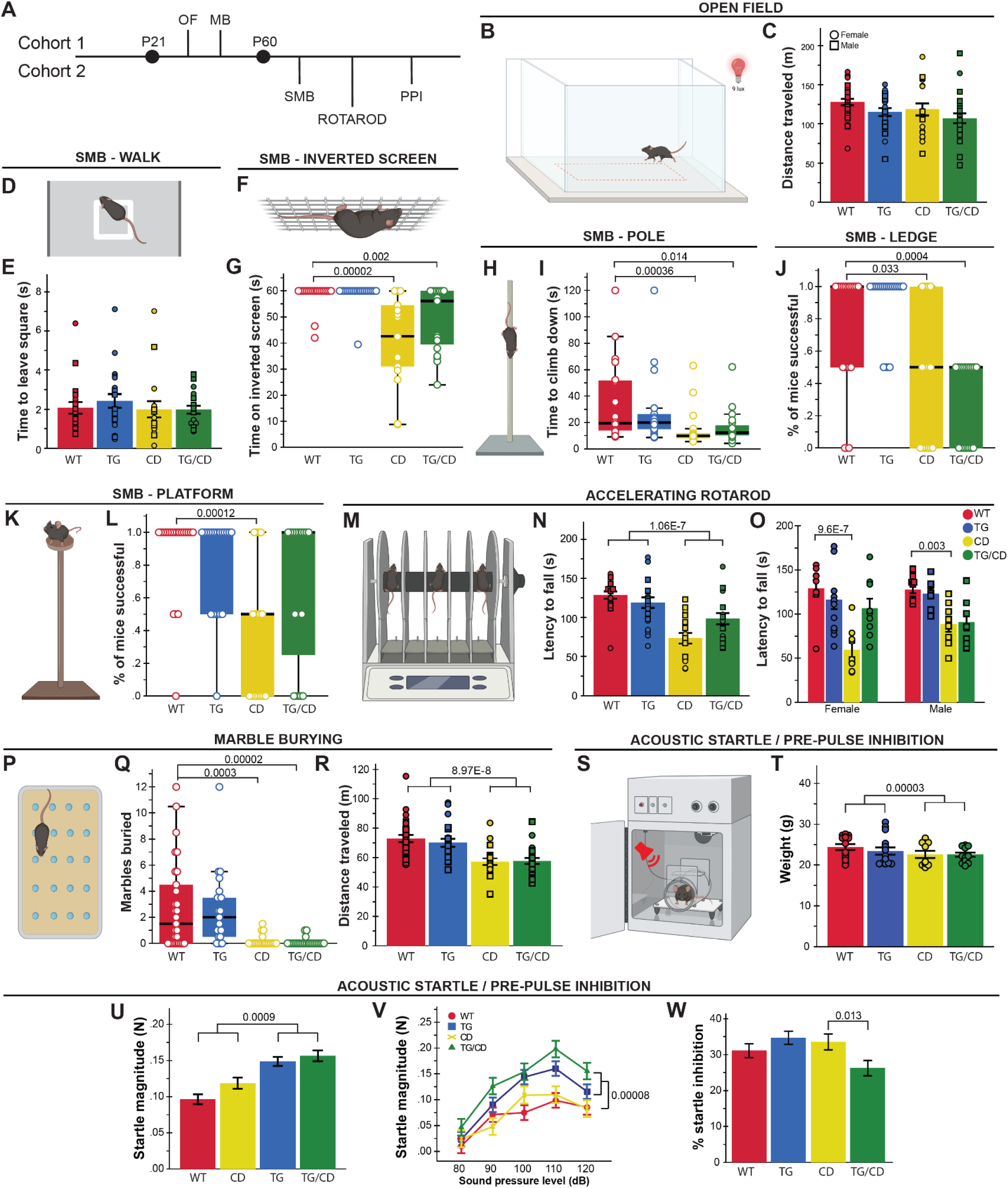
*Gtf2ird1* restoration affects a subset of sensorimotor deficits observed in the CD mice. **A)** Tasks were split between two cohorts; OF = Open Field, MB = Marble Burying, SMB = sensorimotor battery, PPI = Pre-Pulse Inhibition. **B)** A 50×50cm arena under red illumination at 9 lux was used for the 1-hour open field task. **C)** No significant differences were observed between groups in total distance traveled in the open field task. **D)** In the “walk” task, the time for mice to exit the white square in the center of a large open space was measured as a proxy for motor initiation. **E)** No differences in motor initiation were observed. **F)** The inverted screen task measures how long a mouse can hold on for up to 60 seconds **G)** CD and TG/CD mice were not able to hold on to the screen for the full minute. **H)** Mice were placed on a textured pole for up to 120 seconds. **I)** CD and TG/CD mice were significantly faster to leave the pole. **J)** CD and TG/CD mice were unable to stay on an acrylic ledge for a full minute. **K)** Time on a small platform was measured. **L)** CD animals were less successful at remaining on the platform, which was ameliorated with the presence of the transgene. **M)** The Rotarod apparatus used to measure ability to stay on a moving rod. **N)** Animals with the CD allele fell off an accelerating rod faster than WT, but presence of the transgene improved the outcome. **O)** This partial rescue was especially clear in females. **P)** Mice have 30 minutes to explore a chamber with 20 evenly spaced marbles. **Q)** Animals with the CD allele (CD and TG/CD animals) buried significantly fewer marbles than WT and TG animals. **R)** CD and TG/CD animals travel less distance overall, reflected here in significantly fewer center entries. **S)** In the Acoustic Startle/Pre-Pulse Inhibition paradigm, mice are exposed to an acoustic stimulus while confined within a sound-attenuated box on a force plate to measure the startle reflex. **T)** Mice with the CD allele weighed significantly less than those without, requiring a covariate analysis (**U-W**, covariate adjusted means shown). **U)** Transgenic expression of *Gtf2ird1* resulted in a greater startle response (in Newtons) to a 120 dB stimulus and **V)** across various sound levels. **W)** When presented with a pre-pulse, TG/CD mice exhibited a reduced inhibition of startled compared to CD mice. All statistical details including sample sizes are reported in Table S2.

At P30, locomotor activity was not significantly different between groups across the 1-hour open field task (**Fig 2B,C**). In the sensorimotor task at P60, there was no difference between groups in motor initiation in a Walk task (**Fig 2D,E**), though differences in strength and balance were observed in other tasks. Specifically, CD animals were unable to hold on to an inverted screen as long (**Fig 2F,G**; H(3)=30.208, p=1.0×10^−6^) but climbed down the pole faster than WT animals (**Fig 2H,I**; H(3)=16.709, p=8.11×10^−4^). A balance deficit was observed in CD animals as fewer animals in this group were able to remain on a thin, acrylic ledge for a full minute (**Fig 2J**; H(3)=29.487, p=1.1×10^−5^) or on a small platform just large enough for the mice to stand atop (**Fig 2K,L**; H(3)=16.919, p=7.34×10^−4^). Interestingly, rescue of *Gtf2ird1* partially restored performance on the Platform task, as TG/CD animals stayed on the platform significantly longer than their CD counterparts (p=0.046). To examine motor coordination more directly, we used the Rotarod task (**Fig 2M**), which revealed another partial rescue (**Fig 2N**; CD*TG interaction: F(1,69)=6.977, p=0.01). While all mice learned the task and generally improved over subsequent trials, CD and CD/TG animals had a shorter latency to fall relative to WT and TG animals (**Fig 2N**; F(1,69)=35.227, p=1.06×10^−7^). The interaction between CD and TG alleles (e.g., rescue) was most apparent in females (**Fig 2O**; Sex*CD*TG: F(1,69)=4.461, p=0.038).

In the Marble Burying task, both CD and CD/TG mice buried far fewer marbles than the WT and TG groups (**Fig 2P,Q**; H(3)=34.458, p=1.586×10^−7^). This finding is confounded by a matching decrease in total distance traveled (**Fig 2R**; F(1,89)=36.953, p=3.0×10^−8^). Thus, the fewer marbles buried may simply be a factor of hypoactivity, though what is causing the hypoactivity here and not in the open field task is not clear. The effect is perhaps enhanced by the novel bedding used in the Marble Burying task, which was not present in the other apparatus.

Sensory sensitivity is another feature of WS that warrants investigation, as WS individuals are more reactive to sounds (Levitin *et al*. 2005; Gothelf *et al*. 2006). In the Acoustic Startle/Pre-Pulse Inhibition (PPI) task (**Fig 2S**), animals are presented with acoustic stimuli designed to induce the startle response in mice. Animals with the CD allele weighed significantly less compared to all other mice (**Fig 2T**; F(1,47)=21.429, p=0.00003), therefore covariate analysis was used to regress weight out of the model. The CD allele alone did not influence response to an acoustic startle stimulus, but mice harboring the TG allele responded with greater startle magnitude force to the 120 dB startle stimulus (**Fig 2U**; F(1,46)=12.521, p=0.0009) as well as to all sound level presentations (**Fig 2V**; F(1,46)=18.9, p=.00008). These data may indicate a unique feature of unbalanced *Gtf2ird1* expression relative to the rest of the WSCR. The PPI trials revealed an interaction between CD and TG alleles on sensorigating ability (F(1,46)=4,772, p=0.034), with the TG/CD animal showing lower percent inhibition than TG mice (**Fig. 2W**; F(1,46)=6.726, p=0.013).

### Restoring *Gtf2ird1* expression in CD mice rescues light-avoidant but not center-avoidant anxiety-like behaviors

Anxiety is another feature common to WS and Dup7, though the specific forms differ. Non-social anxiety and increased prevalence of phobias are over-represented in the WS population, while Dup7 is characterized by greater social anxiety and separation anxiety, with no clear phenotype related to fear. As there are no specialized treatments for these symptoms among patients, having a well characterized model for preclinical screening of therapeutics may eventually lead to better care. Thus, we thoroughly assessed non-social anxiety-like avoidance features in the CD mouse model to identify tasks sensitive to this mutation and evaluate the potential impact of *Gtf2ird1*.

Anxiety-like behavior is measured in rodents by quantifying passive avoidance behavior in low-threat situations in which perceived danger is diffuse and uncertain (La-Vu *et al*. 2020). These situations for a rodent include open spaces and brightly lit spaces. One common trigger for passive avoidance behavior in rodents is the center space of an open field (**Fig 2B**). Similarly, the Elevated Plus Maze (EPM; **Fig 3D**) measures an animal’s passive avoidance of the open arms. In contrast, light and open space is leveraged together in the Light/Dark Box task, and a decrease in time in the light side of the box has been used to indicate passive avoidance behavior, rather than relying on avoidance of the open space alone. Together, these tasks should inform us of the anxiety-like, passive avoidance features of the CD mice and whether *Gtf2ird1* expression has any effect on these behaviors. Like the sensorimotor tasks, these anxiety-relevant tasks were also split among two cohorts; the 1-hr Open Field and EPM tasks were performed with animals in the first cohort (**Fig 3A**, *above midline*), and the Light/Dark Box task was performed utilizing the second cohort of animals (**Fig 3A**, *below midline*).

**Figure 3.**
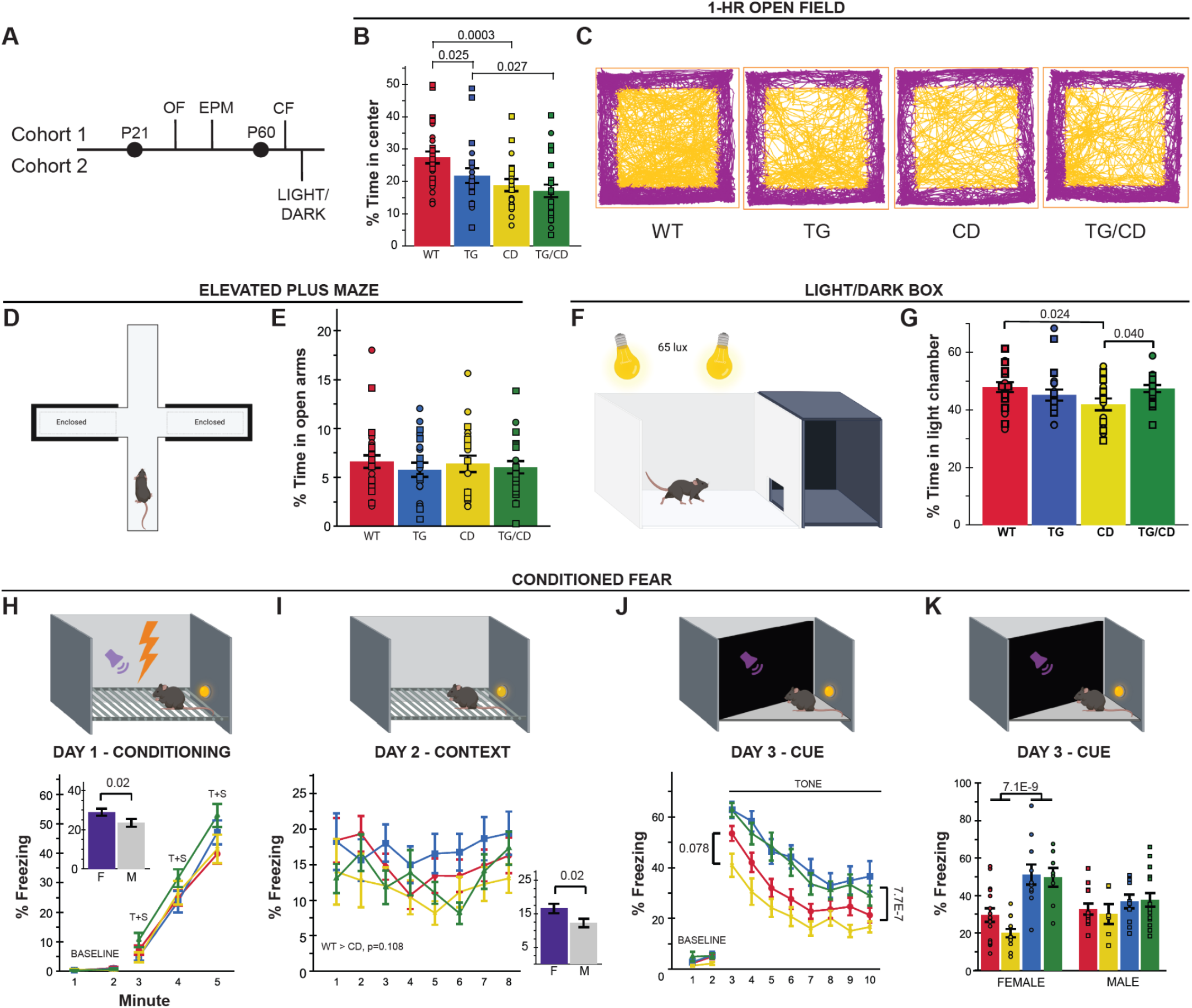
*Gtf2ird1* corrects CD-induced decreased time in light but not in center space. **A)** Tasks related to anxiety and fear were split between two cohorts; OF = Open Field, EPM = Elevated Plus Maze, CF = Conditioned Fear, and LIGHT/DARK = Light/Dark Box. **B)** TG and CD allele decrease time spent in the center of an open field. **C)** Representative track plots of open field task, chosen based on group mean. **D)** Diagram of elevated plus maze apparatus. **E)** No difference in percent time spent in the open arms of the EPM was observed. **F)** Diagram of the Light/Dark Box task. **G)** Decreased time in light side caused by CD allele is rescued with expression the *Gtf2ird1* transgene (TG). **H-K)** *upper panel*, graphic representation of the chamber used for each day of the conditioned fear paradigm, tone is represented by the purple sound icon and shock is represented with the lightning bolt icon. **H)** No significant differences in freezing were observed during the training day of the Conditioned Fear task, though in general females froze more than males (inset). **I)** Day 2 of the Conditioned Fear task measuring context-based fear recall also revealed no significant differences, except between the sexes (inset). **J)** While differences between WT and CD animals were not significant, the TG allele increases percent freezing during cued recall in the third day of Conditioned Fear relative to those animals without that allele. **K)** The TG effect on percent time freezing during cued recall is greater in female mice. T+S = Tone + Shock. All statistical details including sample sizes are reported in Table S3.

In the 1-hr Open Field task conducted under red light at 9 lux, both CD and TG alleles resulted in a decrease in center time (**Fig 3B,C**; CD: F(1,89)=13.956, p=0.0003; TG: F(1,89)=6.862, p=0.010). As the overall distance traveled was not different between groups (**Fig 2C**), these results are consistent with heightened anxiety-like behavior. Regardless of genotype, females spent less time in the center than their male counterparts (Females *M*=18.77, *SD*=7.9; Males *M*=24.8, *SD*=11.7; F(1,89)=14.339, p=0.0003). In contrast to the avoidance behaviors observed in the open field, there were no observed differences in the percent time spent in the open arms of the EPM under complete darkness (**Fig 3D,E**; CD: F(1,86)=0.03, p=0.864; TG: F(1,86)=0.686, p=0.41).

Interestingly, during the Light/Dark Box task (**Fig 3F**), we observed a significant interaction of CD and TG alleles on the percent time spent in the light (**Fig 3G**; F(1,72)=5.250, p=0.025). CD animals spent significantly less time in the light relative to their WT (p=0.024) and TG/CD (p=0.040) counterparts, while TG/CD animals were not significantly different from the WT group, reflective of the TG allele rescuing CD deficits in this task. Thus, *Gtf2ird1* complements the CD mutation for this phenotype.

In contrast to the passive avoidance of anxiety-like measures, fear responses, which have a component of anxiety, are active avoidance behaviors quantified in situations where a threat is imminent and well-defined (La-Vu *et al*. 2020). We used the fear conditioning task in the first cohort of animals to further evaluate active avoidance and associative memory by quantifying freezing behavior in response to a shock paired with a novel auditory cue and spatial context. No differences in freezing response to the pairing of the shock and tone+context were observed between genotypes on Day 1 (**Fig 3H**). However, females froze more than males overall (Day 1, min 3-5; F(1,85)=5.606, p=0.02). This sex effect was also observed during contextual fear recall on Day 2 when mice were re-exposed to the spatial context to test hippocampal-dependent spatial conditioning (**Fig 3I**; F(1,85)=5.650, p=0.02). The CD mice also showed reduced freezing, similar to our previous reports (**Fig 3I,J**) (Nygaard *et al*. 2022), but did not pass the significance threshold.

During amygdalar-dependent cued fear recall on Day 3, animals with the TG allele overexpressing *Gtf2ird1* showed increased freezing in response to the auditory cue (**Fig 3J**; F(1,85)=28.497, p=7.7×10-7). This increased freezing was especially pronounced in females with the TG allele (**Fig 3K**; F(1,85)=11.876, p=0.0009). Mice with only the CD allele showed reduced freezing behavior compared to all other groups, replicating our previous effect (Nygaard *et al*. 2022), although the comparison to WT mice did not pass the significance threshold (p=0.078). Shock sensitivity was comparable across groups (Table S3).

### Enhanced social approach and motivation is independent of *Gtf2ird1*

Finally, given the interesting contrasting social motivation phenotypes in WS and Dup7 patients (Doyle *et al*. 2004; Berg *et al*. 2007), we conducted a comprehensive phenotyping of social behavior in our cohorts (**Fig 4A, 5A**). To identify early signs of social behavior changes, we assessed social communication in pups via the maternal isolation-induced ultrasonic vocalization (USV) paradigm in independent cohort 3 (**Fig 4A,B**). Given elevated aggression in Dup7 patients (Klein-Tasman *et al*. 2022), we included standard measures of social dominance (Tube Test) and aggression (Resident Intruder) measured in cohort 2. Sociability differences in the CD model was originally identified in a modified single chamber version of social approach (Open Field Social Approach), rather than the typical 3-Chamber Social Approach task widely used in ASD models (Crawley 2004; Segura-Puimedon *et al*. 2014). Previous work in our lab failed to identify differences in the 3-Chamber task alone, though on a C57BL/6J x FVB hybrid background that showed lower social approach in general (Kopp *et al*. 2019; Nygaard *et al*. 2019). Thus, our comprehensive battery here included a deliberate precise replication of the Open Field Social Approach conditions as a baseline control in cohort 1 (Sakurai *et al*. 2011; Segura-Puimedon *et al*. 2014), the standard 3-Chamber Social Approach assay in cohort 2 (Crawley 2004; Moy *et al*. 2004), and finally a 14-day Social Operant task we recently designed to be a direct measure of social motivation in rodents in cohort 1 (Chen *et al*. 2021; Maloney *et al*. 2022).

**Figure 4.**
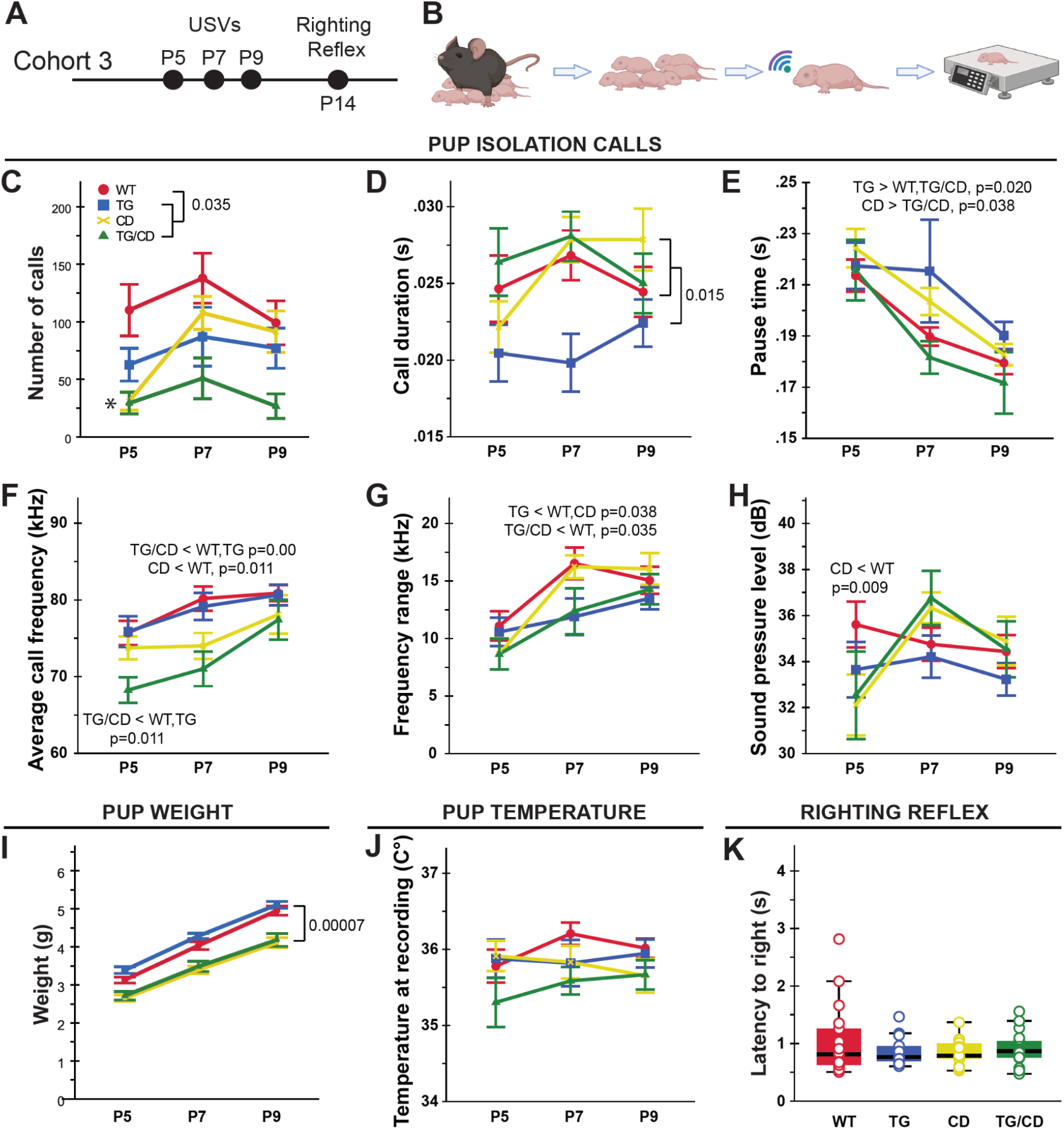
WSCR haploinsufficiency disrupts early communicative behavior. **A)** Cohort 3 was used to assess USVs at P5, 7, and 9, and the righting reflex at P14. **B)** Maternal separation induced pup USV workflow schematic: Remove dam, place pups in nest in warming chamber, individually measure pup temperature while in the nest, record USVs, then weigh and return to nest. **C)** Call rate was reduced in all groups compared to WTs overall (p=0.035). **D-E)** TG mice produced shorter duration calls across all three days, (**D**) with longer pauses between calls on P7 (**E**; TG > WT, TG/CD, p=0.020; CD > TG/CD, p=0.038). **F)** The mean pitch frequency of calls was lower on P5 and P7 for the TG/CD mice (p=0.012) and only P7 for the CD mice (p=0.011). **G)** The range of call frequencies was narrower at P7 for both the TG/CD and TG pups (p=0.038). **H)** At P5, the CD mice were unable to produce calls with similar sound pressure levels to WT pups (p=0.009). **I)** Pup weight after USV recordings show decreased weight in all pups with the CD allele (p=0.00007). **J)** Body temperature at recordings were comparable across groups. **K)** No differences were observed in latency to righting at P14 between groups. All statistical details including sample sizes are reported in Table S3.

Early differences in communication were evident across the three days of the USV task. All groups exhibited a decreased number of calls overall compared to WTs (F(3,83)=7.635, p=.00014; **Fig 4C**). Examination of the spectrotemporal features of the calls revealed TG mice produced overall shorter calls compared to all other groups (F(3,91)=3.411, p=0.020; **Fig 4D**) with increased pause time at P7 (F(3,204)=3.332, p=0.021; **Fig 4E**). Call pitch and sound pressure levels were examined to identify possible laryngeal muscle abnormalities. Mean pitch frequency was lower in the TG/CD pups on P5, and both TG/CD and CD pups on P7 (F(8,129)=4.998, p=0.00002; **Fig 4F**), while the frequency range was narrower on P7 for TG/CD and TG pups (F(8,144)=6.211, p=6.9×10-7; **Fig 4G**). Finally, the sound pressure level, or volume, of calls was lower for CD pups on P5 compared to WTs (F(8,131)=2.742, p=0.008; **Fig 4H**). The differences in USV number do not appear to be due to gross developmental delays as they do not follow the same pattern as the weight data. Specifically, weights were comparable between CD and TG/CD animals from P5 to P9, despite both groups weighing significantly less than WT and TG mice (Geno: F(3,79)=13.619, p=2.8×10-7; **Fig 4I**). No significant differences were observed across groups for pup body temperature during call recordings (**Fig 4J**). Acquisition of the surface righting reflex and eye opening was also assessed at P14. All pups had the ability to flip themselves upright, with no difference in time to exhibit the righting reflex (**Fig 4K**), and no differences in the number of pups with eyes open across groups. Altogether, these data suggest there is an early disruption to social communicative behavior with haploinsufficiency for the WSCR, as well as overexpression of *Gtf2ird1*, which may be driven in part by musculature issues possibly due to loss of the *Eln* (Elastin) gene as suggested by the spectrotemporal call features.

In previous research, adult CD mice on the FVB x C57BL/6J hybrid background showed decreased dominance in the Tube Test and reduced aggression in the Resident Intruder paradigms. In our current study, CD mice on the pure C57BL/6J background seem to win less although no statistically significant difference was observed in the 1 day Tube Test paradigm (**Fig 5B**; H(7)=11.102, p=0.134). In addition, there were no significant differences in the number of attacks exhibited across groups in the Resident Intruder task (**Fig 5C**; CD: F(1,32)=1.167, p=0.288). Thus, the CD mice on the C57BL/6J background failed to exhibit altered hierarchical or agonistic behaviors.

**Figure 5.**
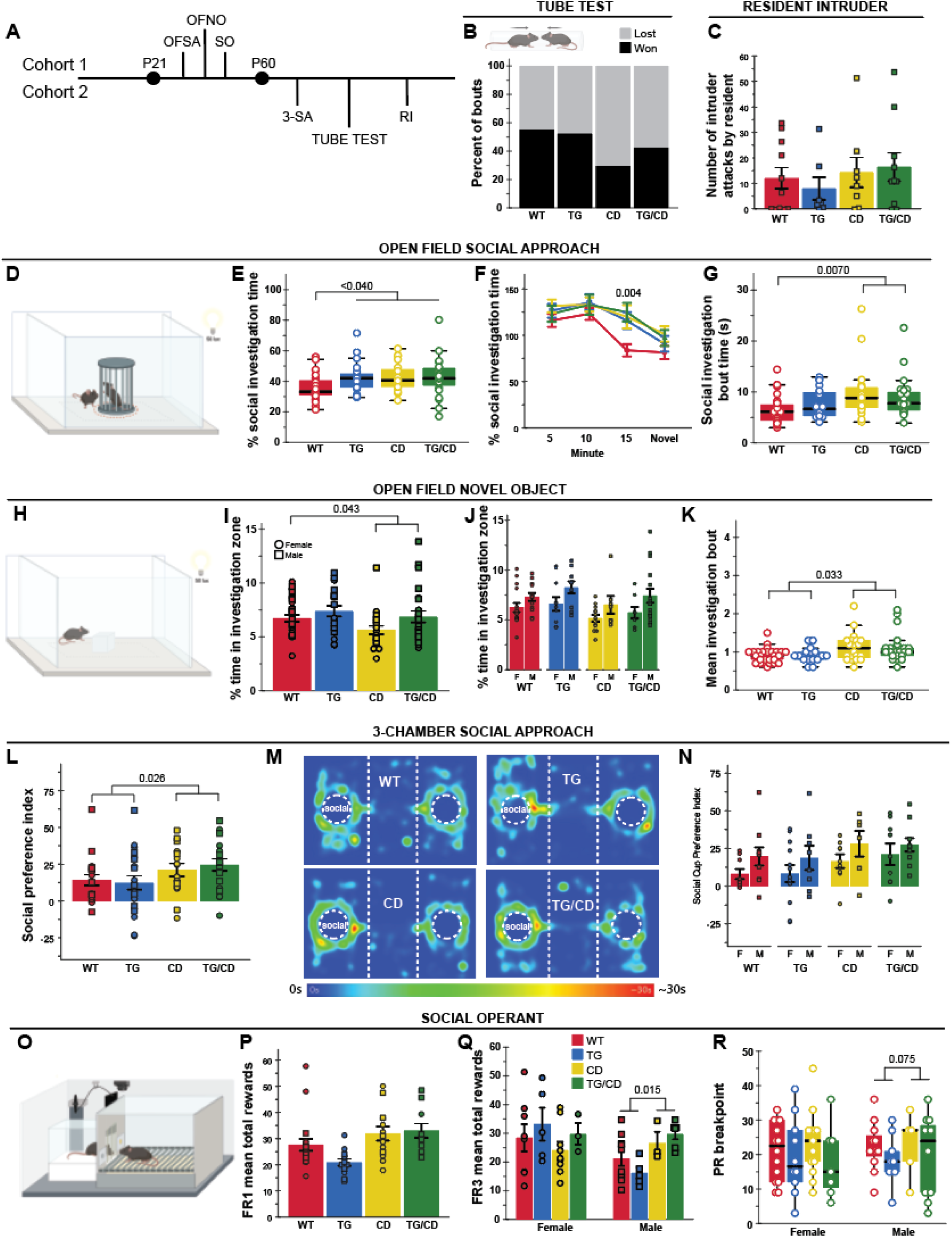
Increased social approach and motivation in the CD model of WS deletion is independent of *Gtf2ird1*. **A)** Tasks related to social behavior were split between two cohorts; OFSA = Open Field Social Approach, OFNO = Open Field Novel Object, SO = Social Operant, 3-SA = 3-Chambered Social Approach, and RI = Resident Intruder. **B)** Apparatus schematic (upper panel). No significant differences measured in the Tube Test for Social Dominance task. **C)** No differences in average number of attacks by resident on intruder. **D)** Open Field Social Approach apparatus schematic. **E)** TG, CD, and TG/CD animals spent a greater percent time investigating a social stimulus. **F)** WT mice showed habituation at 15 minutes that is not seen in the other groups. **G)** Only mice with the CD allele show longer mean bouts of investigation. **H)** Representation of the Open Field Novel Object apparatus. **I)** Novel object avoidance seen in CD and TG/CD animals. **J)** Females in all groups spent less time investigating the object than males. **K)** CD allele caused increased mean investigation bout time in the novel object task. **L)** Social preference index in the 3-chamber social approach task was greater in animals harboring the CD allele. **M)** Representative heatmaps of each group in the 3-chambered social approach task. **N)** Male animals had higher social preference for the social cup than females. **O)** Social Motivation Operant apparatus. **P)** CD allele results in greater mean rewards during FR1. **Q)** Males with the CD allele had increased mean rewards during FR3. **R)** The breakpoint during PR3 was greater in CD and TG/CD animals. circle = female, square = male. All statistical details including sample sizes are reported in Table S4.

Similar to Segura-Puimedon *et al*. (2014), in the Open Field Social Approach task (**Fig 5D**) we found CD animals spent a greater portion of time investigating the social stimulus mouse relative to WT animals (**Fig 5E**; p=0.013). In fact, we observed this increased social approach behavior in all groups relative to WT levels (**Fig 5E**, H(3)=8.916, p=0.03). The significance of this difference appeared to be driven by the lack of habituation, as WT levels of approach fell after 10 minutes while the other groups remained more interested in the social stimulus (**Fig 5F**, H(3)=13.160, p=0.004). While all groups showed sustained interest in the stimulus, the mean social investigation bout time was only higher in CD and TG/CD animals compared to WTs (**Fig 5G**, H(3)=13.16, p=0.006). Regardless, it is clear that the deletion of the WSCR increases social approach behaviors here as measured in investigation time and average investigation bout. Further, these findings replicate the previously observed approach behavior in the modified social approach task (Segura-Puimedon *et al*. 2014).

To control for the potential impact of novelty on the Open Field Social Approach task (**Fig 5D**), we next tested the reaction to a non-social novel object, an acrylic square placed in the center of the familiar open field arena (**Fig 5H**). Animals with the CD allele spent significantly less time near the novel object compared to WT mice (**Fig 5I**; F(1,86)=4.203, p=0.043). TG allele alone did not have an effect. In addition to the effect of the CD allele, females of all groups spent less time investigating the novel object as well (**Fig 5J**; F(1,86)=11.901, p=8.71×10^−4^), possibly due to increased exploration of the chamber. Though less time overall was spent investigating the object, mean investigation bout time was higher in animals with the CD deletion, just as it was in the Open Field Social Approach task (**Fig 5K**; H(3)=11.271, p=0.01). These data suggest that while some features of investigation may be shared across social and non-social stimulus, such as investigation bout time, other features such as total time are dependent on the valence of the stimuli, with the CD allele increasing interest in a social stimulus but decreasing interest in a non-social novel object.

The increased social approach in CD mice was also observed in the classic 3-Chamber Social Approach task. Deleting the WS region results in a greater preference for the social cup, compared to an empty cup (**Fig 5L,M**; F(1,66)=4.990, p=0.029). This preference was especially pronounced in males across all genotypes (**Fig 5N**; F(1,66)=5.414, p=0.023). No differences were observed between alleles for social novelty preference (**Table 5**).

Finally, we applied our social operant task to precisely investigate how social motivation may fluctuate with the loss of the WSCR (**Fig 5O**). Among the animals that showed conditioning for the social reward, those with the complete deletion allele (CD and CD/TG genotypes), reached a higher number of total rewards during FR1 (**Fig 5P**; F(1,47)=14.07, p=4.83×10^−4^). During FR3, an interaction between sex and CD allele emerged, showing that only males with the CD allele reach more rewards (**Fig 5Q**; CD*Sex: F(1,44)=4.932, p=0.032). In addition, males with the CD allele resulted in a higher breakpoint during PR3 (**Fig 5R**; H(3)=8.092, p=0.044). These results show that complete deletion of the WSCR in mice resulted in higher social motivation, meaning they were willing to work harder to keep accessing the social stimuli. While this appears to be driven by an increase in males, the limited number of animals who met criteria to be considered learners (and thus be included in our analysis) constrains our power to assess sex effects.

Williams Syndrome is characterized by high anxiety, yet in non-social domains. Thus, we leveraged several of our related tasks to assess open area avoidance behaviors in the CD mice in both social and non-social settings. The mice with the CD allele spent a smaller percent of time in the center of the apparatus in both the 1-hr Open Field task (**Fig 6B,C**; F(1,86)=11.930, p=0.0009) and the Open Field Novel Object task (**Fig 6D,E**; F(1,86)=6.070, p=0.016). We also observed this center avoidance by the CD mice in the Marble Burying task (**Fig 6F,G**; F(1,86)=19.826, p=0.00003). However, in the Open Field Social Approach task, the CD mice spent as much time in the center as all other groups (**Fig 6H,I**; F(1,85)=3.336, p=0.071). Thus, these observations are consistent with the dichotomous anxiety profile of WS, defined by prevalent anxiety that does not present as social anxiety.

**Figure 6.**
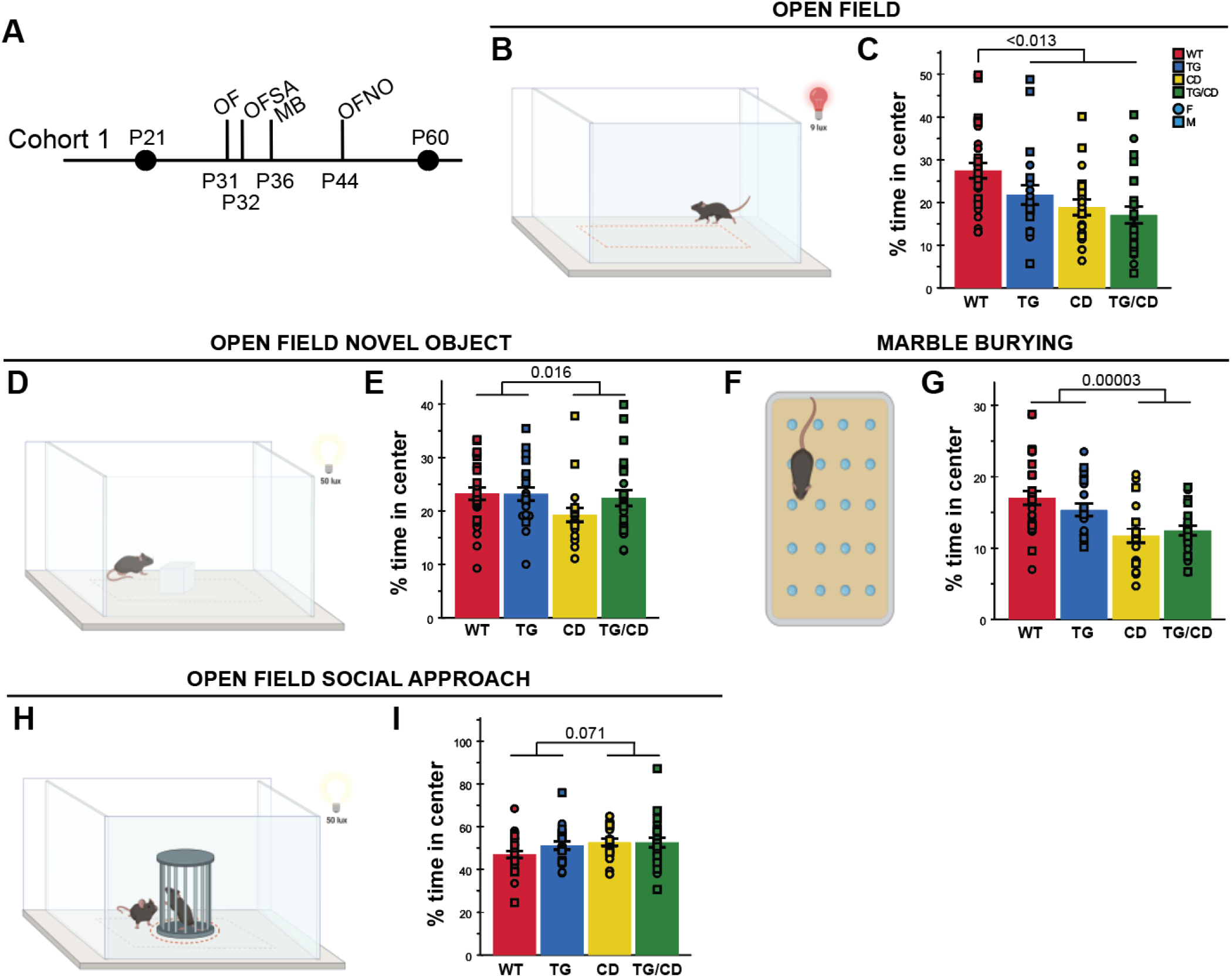
CD mice exhibited increased open area avoidance behavior only in non-social settings. **A)** Tasks allowing for center avoidance quantification in Cohort 1; OF = Open Field, OFSA = Open Field Social Approach, MB = Marble Burying, and OFNO = Open Field Novel Object. **B)** Open Field apparatus schematic. **C)** TG, CD, and TG/CD mice spent less percent of time in the center. **D)** Open Field Novel Object apparatus schematic. **E)** CD and TG/CD animals spent a lesser percent time in the center of the apparatus. **F)** Marble Burying apparatus schematic. **G)** CD and TG/CD animals spent a lesser percent time in the center of the Marble Burying apparatus. **H)** Representation of the Open Field Social Approach apparatus. **I)** Comparable percentage of time was spent in the apparatus center for all groups. circle = female, square = male. All statistical details including sample sizes are reported in Table S4.

## Discussion

In this extensive characterization of mouse models relevant to the WSCR, the consequences of complete deletion of the region were obvious. We found a widespread effect of WSCR deletion in the CD model, causing deficits in sensorimotor abilities (i.e., performance in Inverted Screen, Ledge, Platform, Rotarod, Marble Burying tasks), select anxiety-like behaviors (i.e., time in Open Field center, and time in light side of the Light/Dark Box), and enhanced social interest (i.e., approach behaviors in open field and 3-chamber set-ups, and increased motivation in the social operant paradigm). Mice with the WSCR deletion did not bury as many marbles though this result was confounded by a strong avoidance of the center; given some of the strength, balance, and coordination issues reported above (Fig 2), it may be that these mice have a harder time navigating the novel aspen bedding. We also presented a novel transgenic line expressing a gene of interest, *Gtf2ird1*, and used this line to genetically rescue *Gtf2ird1* expression in the CD line to examine its ability to rescue any of the atypical phenotypes presented. The transgenic *Gtf2ird1* line highlighted a role for *Gtf2ird1* in sensorimotor coordination (as evidenced in the Platform and Rotarod tasks) and potentially sensory processing more generally (sound sensitivity and potentially light sensitivity). While *Gtf2ird1* affected a few features clearly, most features were not significantly impacted by its rescue or overexpression, suggesting either *Gtf2ird1* is not involved or is only part of the underlying etiology and would require rescuing multiple genes in parallel to see an effect.

Interestingly, *Gtf2ird1* seemed linked with a hyper-response to sound as demonstrated by force produced during acoustic startle trials during PPI, and increased freezing during contextual fear; this effect was consistent regardless of the presence of the CD allele This could mean one of two things: either that the HA-tagged beta isoforms have an altered function specifically affecting these phenotypes, or that any discrepancy in *Gtf2ird1* expression relative to the other genes in the WSCR impacts these phenotypes. Either possibility is potentially interesting. If it is an issue of the HA tag, we learn that a specific subset of *Gtf2ird1’s* isoforms is involved in responsivity to sound, or if it’s an issue of discordant expression within the WSCR, we learn there is an interaction between *Gtf2ird1* and at least one other gene in the region. Unfortunately, distinguishing these possibilities would require the generation of an additional, untagged, *Gtf2ird1* BAC transgenic line, which is beyond the scope of the current study.

Beyond its usefulness here, the novel transgenic *Gtf2ird1* line may also provide opportunities to research into the various roles *Gtf2ird1* isoforms may play in typical development. *Gtf2ird1* is an extensively alternatively spliced gene, with numerous uncharacterized isoforms. Our TG-Gtf2ird1-HA mouse tags less than half of the isoforms (only the beta variants that contain the full exon 30) (Tay *et al*. 2003); leveraging this fact, we can investigate how these two groups of isoforms differ. Utilizing an HA antibody in a pull-down assay would effectively separate these isoform groups for downstream analysis to compare their functions, particularly in regard to genomic binding.

Synthesizing the results of this study beyond the contributions of *Gtf2ird1*, we show the Complete Deletion mouse showed an interesting variety of responses when assessed for anxiety-related phenotypes, given that non-social phobias and anxiety are important components of the Williams Syndrome deletion in people. Avoidance of the center in the Open Field task suggested an anxiety-like phenotype, which was replicated in the decreased time in the light side of the Light/Dark Box, but no effect was apparent in the EPM. It may be that the EPM is not sensitive to the particular anxiety-inducing features relevant to WS. It is possible that the differences observed in the Open Field and EPM tasks (Fig 3) may be due to differences in the length of the tasks (60 vs 5 min, respectively), or that the limited time animals spent in the EPM open arms (<10% of time on average) results in reduced sensitivity to detect group differences with small effects sizes. Despite inconsistency among these tasks, center avoidance was also present in tasks not typically analyzed for center time (Fig. 6; Open Field Novel Object and Marble Burying), showing that aspect of anxiety-related behavior is consistent across tasks. The only exception was the Open Field Social Approach task, discussed more below. Thus, future studies to understand the circuits mediating the partial anxiety-like features in CD mice should focus on Open Field and Light/Dark box. Such studies focusing on mechanism discovery for behavioral symptoms could be a way to provide meaningful answers despite complicated etiology, as suggested by Kozel *et al*. (2021).

The hypersocial phenotype that has been documented in people with WS is also recapitulated in the CD mice in multiple behavioral tasks (Open Field Social Approach, 3-Chamber Social Approach, and Social Motivation Operant Conditioning). Additionally, while an avoidance of the center space was consistently seen across many tasks, it was not observed in the Open Field Social Approach task, showing how the presence of a conspecific can potentially overcome the center anxiety seen when no social stimulus is present. Whether this means social stimuli can be more salient than anxiety cues or whether this indicates a separate circuit for social anxiety, the CD mouse model is appropriate for further teasing apart the underpinnings of this hypersocial phenotype as well as the anxiety-like features mentioned above.

## Acknowledgments

The authors would like to thank Amanda Titus and the Animal Behavior Subunit of the IDDRC at Washington University School of Medicine for assistance in running select assays, Carly Wender and Gunnar Forsberg for validation of the *Gtf2ird1* overexpressing mice, Dr. Victoria Campuzano for sharing the CD mice, Nastacia Goodwin from the University of Washington - Seattle for technical support for the SimBA program, and the transgenics core at UC Irvine for production of the TG mice. Some elements of the figures were created using BioRender.com. This work was supported by the NSF (DGE-1745038 to KRN) and the NIMH (MH094604 (JV), R01MH067234 (KM), R01MH107515 (JDD)), and NICHD P50HD103525 (IDDRC@WUSTL).

## Data Availability Statement

All data and detailed protocols are available upon reasonable request.

## Supplementary Tables

**Table S1. Statistical information for Figure 1 – Molecular Validation. Table S2. Statistical information for Figure 2 – Sensorimotor Tasks**

**Table S3. Statistical information for Figures 3 and 4 – Anxiety and Fear-Related Tasks, and Developmental Assessment**

**Table S4. Statistical information for Figures 5 and 6 – Social Behavior Tasks and Center Avoidance Assessment**

## Notes

### Competing Interest Statement

The authors have declared no competing interest.

## References

Adamo, A., Atashpaz, S., Germain, P.-L., Zanella, M., D’Agostino, G., Albertin, V., Chenoweth, J., Micale, L., Fusco, C., Unger, C., Augello, B., Palumbo, O., Hamilton, B., Carella, M., Donti, E., Pruneri, G., Selicorni, A., Biamino, E., Prontera, P., McKay, R., Merla, G. & Testa, G. (2014) 7q11.23 dosage-dependent dysregulation in human pluripotent stem cells affects transcriptional programs in disease-relevant lineages. Nature Genetics 47, ng.3169.

Berg, J.S., Brunetti-Pierri, N., Peters, S.U., Kang, S.-H.L., Fong, C.-T., Salamone, J., Freedenberg, D., Hannig, V.L., Prock, L.A., Miller, D.T., Raffalli, P., Harris, D.J., Erickson, R.P., Cunniff, C., Clark, G.D., Blazo, M.A., Peiffer, D.A., Gunderson, K.L., Sahoo, T., Patel, A., Lupski, J.R., Beaudet, A.L. & Cheung, S.W. (2007) Speech delay and autism spectrum behaviors are frequently associated with duplication of the 7q11.23 Williams-Beuren syndrome region. Genet Med 9, 427–441.

Chen, J., Lambo, M.E., Ge, X., Dearborn, J.T., Liu, Y., McCullough, K.B., Swift, R.G., Tabachnick, D.R., Tian, L., Noguchi, K., Garbow, J.R., Constantino, J.N., Gabel, H.W., Hengen, K.B., Maloney, S.E. & Dougherty, J.D. (2021) A MYT1L syndrome mouse model recapitulates patient phenotypes and reveals altered brain development due to disrupted neuronal maturation. Neuron 109, 3775–3792.e14.

Chevallier, C., Kohls, G., Troiani, V., Brodkin, E.S. & Schultz, R.T. (2012) The social motivation theory of autism. Trends in Cognitive Sciences 16, 231–239.

Crawley, J.N. (2004) Designing mouse behavioral tasks relevant to autistic-like behaviors. Mental Retardation and Developmental Disabilities Research Reviews 10, 248–258.

Doyle, T.F., Bellugi, U., Korenberg, J.R. & Graham, J. (2004) “Everybody in the world is my friend” hypersociability in young children with Williams syndrome. Am J Med Genet A 124A, 263–273.

Ebert, P.J., Campbell, D.B. & Levitt, P. (2006) Bacterial artificial chromosome transgenic analysis of dynamic expression patterns of regulator of G-protein signaling 4 during development. I. Cerebral cortex. Neuroscience 142, 1145–1161.

Gothelf, D., Farber, N., Raveh, E., Apter, A. & Attias, J. (2006) Hyperacusis in Williams syndrome: characteristics and associated neuroaudiologic abnormalities. Neurology 66, 390–395.

Haridas, S., Ganapathi, R., Kumar, M. & Manda, K. (2018) Whisker dependent responsiveness of C57BL/6J mice to different behavioral test paradigms. Behavioural Brain Research 336, 51–58.

Klein-Tasman, B.P., Yund, B.D. & Mervis, C.B. (2022) The Behavioral Phenotype of 7q11.23 Duplication Syndrome Includes Risk for Oppositional Behavior and Aggression. Journal of Developmental & Behavioral Pediatrics 10.1097/DBP.0000000000001068.

Kopp, N., McCullough, K., Maloney, S.E. & Dougherty, J.D. (2019) Gtf2i and Gtf2ird1 mutation do not account for the full phenotypic effect of the Williams syndrome critical region in mouse models. Hum Mol Genet.

Kopp, N.D., Nygaard, K.R., Liu, Y., McCullough, K.B., Maloney, S.E., Gabel, H.W. & Dougherty, J.D. (2020) Functions of Gtf2i and Gtf2ird1 in the developing brain: transcription, DNA binding and long-term behavioral consequences. Hum Mol Genet 29, 1498–1519.

Kozel, B.A., Barak, B., Kim, C.A., Mervis, C.B., Osborne, L.R., Porter, M. & Pober, B.R. (2021) Williams syndrome. Nat Rev Dis Primers 7, 1–22.

La-Vu, M., Tobias, B.C., Schuette, P.J. & Adhikari, A. (2020) To Approach or Avoid: An Introductory Overview of the Study of Anxiety Using Rodent Assays. Front Behav Neurosci 14, 145.

Levitin, D.J., Cole, K., Lincoln, A. & Bellugi, U. (2005) Aversion, awareness, and attraction: investigating claims of hyperacusis in the Williams syndrome phenotype. Journal of Child Psychology and Psychiatry 46, 514–523.

Maloney, S.E., Akula, S., Rieger, M.A., McCullough, K.B., Chandler, K., Corbett, A.M., McGowin, A.E. & Dougherty, J.D. (2018a) Examining the reversibility of long-term behavioral disruptions in progeny of maternal SSRI exposure. Eneuro 5.

Maloney, S.E., Chandler, K.C., Anastasaki, C., Rieger, M.A., Gutmann, D.H. & Dougherty, J.D. (2018b) Characterization of early communicative behavior in mouse models of neurofibromatosis type 1. Autism Research 11, 44–58.

Maloney, S.E., Sarafinovska, S., Weichselbaum, C., McCullough, K.B., Swift, R.G., Liu, Y. & Dougherty, J.D. (2022) A comprehensive assay of social motivation reveals sex-differential roles of ASC-associated genes and oxytocin.

Maloney, S.E., Yuede, C.M., Creeley, C.E., Williams, S.L., Huffman, J.N., Taylor, G.T., Noguchi, K.N. & Wozniak, D.F. (2019) Repeated neonatal isoflurane exposures in the mouse induce apoptotic degenerative changes in the brain and relatively mild long-term behavioral deficits. Sci Rep 9, 2779.

Morris, C.A., Mervis, C.B., Paciorkowski, A.P., Abdul-Rahman, O., Dugan, S.L., Rope, A.F., Bader, P., Hendon, L.G., Velleman, S.L., Klein-Tasman, B.P. & Osborne, L.R. (2015) 7q11.23 Duplication Syndrome: Physical Characteristics and Natural History. Am J Med Genet A 167A, 2916–2935.

Moy, S.S., Nadler, J.J., Perez, A., Barbaro, R.P., Johns, J.M., Magnuson, T.R., Piven, J. & Crawley, J.N. (2004) Sociability and preference for social novelty in five inbred strains: an approach to assess autistic-like behavior in mice. Genes, Brain and Behavior 3, 287–302.

Nath, T., Mathis, A., Chen, A.C., Patel, A., Bethge, M. & Mathis, M.W. (2019) Using DeepLabCut for 3D markerless pose estimation across species and behaviors. Nat Protoc 14, 2152–2176.

Nilsson, S.R., Goodwin, N.L., Choong, J.J., Hwang, S., Wright, H.R., Norville, Z.C., Tong, X., Lin, D., Bentzley, B.S., Eshel, N., McLaughlin, R.J. & Golden, S.A. (2020) Simple Behavioral Analysis (SimBA) – an open source toolkit for computer classification of complex social behaviors in experimental animals.

Nygaard, K.R., Maloney, S.E. & Dougherty, J.D. (2019) Erroneous inference based on a lack of preference within one group: Autism, mice, and the social approach task. Autism Research 12, 1171–1183.

Nygaard, K.R., Swift, R.G., Glick, R.M., Wagner, R.E., Maloney, S.E., Gould, G.G. & Dougherty, J.D. (2022) Oxytocin receptor activation does not mediate associative fear deficits in a Williams Syndrome model. Genes, Brain and Behavior 21, e12750.

Ramos, A. (2008) Animal models of anxiety: do I need multiple tests? Trends in Pharmacological Sciences 29, 493–498.

Sakurai, T., Dorr, N.P., Takahashi, N., McInnes, L.A., Elder, G.A. & Buxbaum, J.D. (2011) Haploinsufficiency of Gtf2i, a gene deleted in Williams Syndrome, leads to increases in social interactions. Autism Res 4, 28–39.

Schneider, T., Skitt, Z., Liu, Y., Deacon, R.M.J., Flint, J., Karmiloff-Smith, A., Rawlins, J.N.P. & Tassabehji, M. (2012) Anxious, hypoactive phenotype combined with motor deficits in Gtf2ird1 null mouse model relevant to Williams syndrome. Behavioural Brain Research 233, 458–473.

Segura-Puimedon, M., Sahún, I., Velot, E., Dubus, P., Borralleras, C., Rodrigues, A.J., Valero, M.C., Valverde, O., Sousa, N., Herault, Y., Dierssen, M., Pérez-Jurado, L.A. & Campuzano, V. (2014) Heterozygous deletion of the Williams–Beuren syndrome critical interval in mice recapitulates most features of the human disorder. Hum Mol Genet 23, 6481–6494.

Strong, E., Butcher, D.T., Singhania, R., Mervis, C.B., Morris, C.A., De Carvalho, D., Weksberg, R. & Osborne, L.R. (2015) Symmetrical Dose-Dependent DNA-Methylation Profiles in Children with Deletion or Duplication of 7q11.23. Am J Hum Genet 97, 216–227.

Tay, E.S.E., Guven, K.L., Subramaniam, N., Polly, P., Issa, L.L., Gunning, P.W. & Hardeman, E.C. (2003) Regulation of alternative splicing of Gtf2ird1 and its impact on slow muscle promoter activity. Biochemical Journal 374, 359–367.

Young, E.J., Lipina, T., Tam, E., Mandel, A., Clapcote, S.J., Bechard, A.R., Chambers, J., Mount, H.T.J., Fletcher, P.J., Roder, J.C. & Osborne, L.R. (2008) Reduced fear and aggression and altered serotonin metabolism in Gtf2ird1-targeted mice. Genes Brain Behav 7, 224–234.

Yuede, C.M., Olney, J.W. & Creeley, C.E. (2013) Developmental Neurotoxicity of Alcohol and Anesthetic Drugs Is Augmented by Co-Exposure to Caffeine. Brain Sciences 3, 1128–1152.

Zanella, M., Vitriolo, A., Andirko, A., Martins, P.T., Sturm, S., O’Rourke, T., Laugsch, M., Malerba, N., Skaros, A., Trattaro, S., Germain, P.-L., Mihailovic, M., Merla, G., Rada-Iglesias, A., Boeckx, C. & Testa, G. (2019) Dosage analysis of the 7q11.23 Williams region identifies BAZ1B as a major human gene patterning the modern human face and underlying self-domestication. Science Advances 5, eaaw7908.

